# Mechanical control of cell proliferation patterns in growing tissues

**DOI:** 10.1101/2023.07.25.550581

**Authors:** Logan C. Carpenter, Fernanda Pérez-Verdugo, Shiladitya Banerjee

## Abstract

Cell proliferation plays a crucial role in regulating tissue homeostasis and development. However, our understanding of how cell proliferation is controlled in densely packed tissues is limited. Here we develop a computational framework to predict the patterns of cell proliferation in growing tissues, connecting single-cell behaviors and cell-cell interactions to tissue-level growth. Our model incorporates probabilistic rules governing cell growth, division, and elimination, while also taking into account their feedback with tissue mechanics. In particular, cell growth is suppressed and apoptosis is enhanced in regions of high cell density. With these rules and model parameters calibrated using experimental data, we predict how tissue confinement influences cell size and proliferation dynamics, and how single-cell physical properties influence the spatiotemporal patterns of tissue growth. Our findings indicate that mechanical feedback between tissue confinement and cell growth leads to enhanced cell proliferation at tissue boundaries, whereas cell growth in the bulk is arrested. By tuning cellular elasticity and contact inhibition of proliferation we can regulate the emergent patterns of cell proliferation, ranging from uniform growth at low contact inhibition to localized growth at higher contact inhibition. Furthermore, mechanical state of the tissue governs the dynamics of tissue growth, with cellular parameters affecting tissue pressure playing a significant role in determining the overall growth rate. Our computational study thus underscores the impact of cell mechanical properties on the spatiotemporal patterns of cell proliferation in growing tissues.

## INTRODUCTION

The regulation of cell proliferation is critical for tissue development, maintenance of homeostasis, and disease suppression (1, 2). Uncontrolled proliferation results in hyperplasia, a hallmark of cancerous mutations (3). Cell proliferation involves the coordinated processes of cell growth and division at the individual cell level. However, the precise coordination of cell growth and division within the multicellular context is still poorly understood. One mechanism implicated in the control of cell growth is contact inhibition of proliferation, where cellular growth is inhibited in denser environments (4). Contact inhibition has been shown to be regulated by several factors, including intercellular signaling (5–7), cellular crowding (8, 9) mechanical stress (10–12) and pressure (13). In particular, cells under compression grow slower than cells under stretch (10, 14). However, cellular response to mechanical stress and intercellular signaling is influenced by various factors, such as cell-cell adhesion, cellular resistance to deformations, cytoskeletal tension, local cell density, and cell cycle phase. Manipulating these factors individually within tissues is challenging, leaving the relative contributions of mechanical and biochemical processes in regulating cell growth largely unexplored.

Cell proliferation relies crucially on the mechanism of cell division. While cell growth and division are coupled in single cells (15, 16), cell division can occur independently of growth in tissues (17). Various models of cell division control have been proposed in recent years (15). These include the *adder model* (18), where cells grow by a fixed volume between successive divisions, and the *sizer model* (19), where cell division occurs above a critical cell size threshold (8, 20). Recent studies, however, suggested that the regulation of the cell cycle in both single cells and tissues can be well described by a *G1 sizer model* (21, 22). In this model, there is a cell size-dependent transition from the growth (G1) phase to the mitosis (S/G2/M) phase. Cells spend a fixed amount of time in S/G2/M phase before division occurs. Consequently, the rate of cell division is regulated by the time required to reach the critical size threshold at the G1/S transition, which, in turn, depends on cellular growth rate. Considering the diverse range of biophysical factors influencing cell size and growth rate, it remains to be determined how cell proliferation patterns in tissues are precisely controlled by these factors.

Many cell-based models of tissue growth have been developed in recent years, which explored the effects of contact inhibition (23–25), cellular crowding (9, 26), mechanical stress (8, 10, 11) and intercellular adhesion (27) on tissue-scale growth dynamics. In particular, it has been observed that contact inhibition of proliferation leads to enhanced cell proliferation at tissue boundaries while inhibiting growth within the bulk of the tissue (8, 10). Notably, experiments have revealed diverse patterns of tissue growth, including bulk growth (22, 28, 29) and spatially patterned growth (28, 30). How the emergent patterns of cell proliferation are regulated by cell-cell mechanical interactions remains unclear.

In this work, we developed a cell-based model for tissue growth that aims to uncover how single-cell behaviors and mechanical properties govern tissue-scale proliferation patterns. Our integrative model couples the physical dynamics of cells with active cellular behaviors that determine their fates (Fig. 1). The physical layer of the model is implemented using the Cellular Potts Model (31), whereas the active behaviors are implemented as rules for cell cycle regulation, contact inhibition of proliferation, and cell elimination via apoptosis and extrusion. In contrast to previous cellular models for tissue growth (8, 10, 25, 26, 32, 33), we incorporate a G1 sizer model for cell cycle progression (22) that allows for tunable growth rates of individual cells based on the amount of crowding. Furthermore, our model implements contact inhibition through crowding-induced suppression of cellular growth rate and introduces a cell density-dependent probability of apoptosis. Taken together, our model incorporates feedback between mechanical pressure and the rates of cell growth, division, and apoptosis within a tissue context.

**Figure 1:**
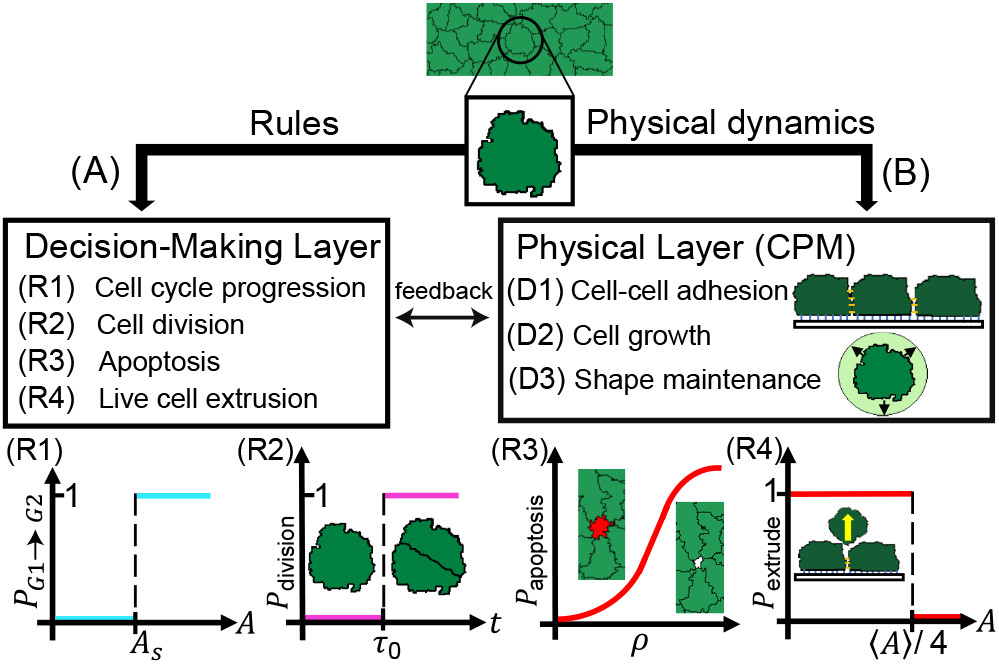
Overview of the computational modeling framework. (A) Cell-autonomous decisions and their rules. (R1) Cycle progression from G1 (sizer-phase) to G2 (timer-phase) phase is triggered by the cell area exceeding a threshold value *As*. (R2) Mitosis is triggered after a time τ ^0^ has elapsed since the beginning of the S/G2/M phase. (R3) Apoptosis is triggered probabilistically as a function of local cell density. (R4) Cellular extrusion is triggered when the cell area reaches one fourth of the mean area for all cells in the tissue. (B) The physical layer evolves via probabilistic minimization of cell mechanical energy using the cellular Potts model that accounts for: (D1) contact energy at cell-cell and cell-medium interfaces, (D2) growth in cell size by increasing cell target area and subsequent relaxation of area elastic energy, and (D3) cell shape maintenance by relaxing mechanical energy of the cell.

Using this model, we successfully recapitulated several single-cell behaviors observed experimentally, including the distribution of cell areas, the control of cell cycle duration, and the decrease in cell size over time. The latter is caused by a transition from a *sizer-like* to a *timer-like* control of cell size that leads to volume-reductive cell division as local cell density is increased via tissue growth. Subsequently, cell cycle arrest occurs as cell size falls below the minimum size threshold required for G1/S transition. Notably, we found that the cell size threshold at the G1/S transition is the primary determinant of tissue properties in homeostasis, including homeostatic density, steady-state rates of cell division and elimination. In contrast, the rate of cell proliferation is predominantly regulated by the physical properties of cells, specifically cell elasticity, growth rate and sensitivity to contact inhibition. Interestingly, spatial patterns of cell growth and cell cycle exhibit variations depending on the values of cell elastic modulus and sensitivity to contact inhibition. Tissues with moderate to high sensitivities to contact inhibition display boundary growth, while tissues with a higher cell elastic modulus exhibit bulk growth. For intermediate values of elastic modulus and contact inhibition, diverse patterns of cell proliferation emerge, emphasizing the influential role of mechanical factors in governing tissue-scale proliferation dynamics. These findings provide new insights into the intricate interplay between mechanical and cellular factors in tissue growth regulation.

## MODEL

To predict the emergent patterns of cell proliferation, we developed a two-dimensional cell-based model for tissue growth that integrates two distinct computational layers – (1) a physical layer that simulates mechanical interactions between cells and between cells and the external medium, (2) a decision-making layer that encodes cell-autonomous rules for growth, division and apoptosis (Fig. 1). For simplicity, we neglected cell motility and focused on the role of cell mechanics and cell cycle regulation on tissue growth.

In the physical layer, we used the lattice-based Cellular Potts model (31, 34), where cells are formed by a subset of lattice sites sharing the same identity. Each lattice site *i* is assigned a type τ_*i*_ = {c: cell, m: medium}, and an integer-valued identity *n*_*i*_, with *n*_*i*_ = 0 for the external medium, and *n*_*i*_ *>* 0 for a cell. The mechanical energy of cells in the Cellular Potts Model can be written in a simplified form as:

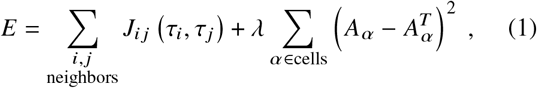

where the first term represents interfacial energy arising from tension at cell-cell junctions, adhesion between cells and between cells and the extracellular medium. The interfacial energy is summed over adjacent lattice-sites *i* and *j*, with *n*_*i*_ ≠*n* _*j*_. The energy *J*_*i j*_ can take two different values depending on the adjacent lattice types: *J*_*i j*_ (c, c) = *J*_cc_, and *J*_*i j*_ (c, m) = *J*_*i j*_ (m, c) = *J*_cm_. We assumed greater adhesion for cell-cell contact than cell-medium contact yielding the condition *J*_*cc*_ ≤*J*_*cm*_. The second energy term is summed over all the cells, representing an elastic energy that penalizes deviations of the cell areas *A*_*α*_ from their preferred (target) values 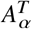, with an area elastic modulus *λ*. At each computational time-step the lattice site *i* can take the value of its adjacent site *n* _*j*_ (with *n*_*i*_ ≠ *n* _*j*_), inducing cell shape changes. These lattice moves are executed via the Metropolis-Hastings algorithm (35, 36) to ensure that the moves minimize the mechanical energy.

The decision-making layer implements cell-autonomous rules for cell cycle progression, mitosis, apoptosis, extrusion, and changing the cell’s target area. Cell growth is implemented by increasing the cell’s target area *A*^*T*^ (26), as

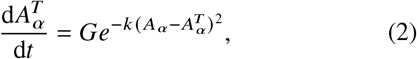

where the constant *G* represents the rate of growth in the absence of crowding (when 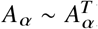). We define *k* as a metric of cellular sensitivity to crowding. In crowded conditions, it is difficult for cells to maintain their areas close to the growing target area, and hence 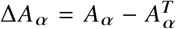 increases in magnitude. Under these conditions, Eq. (2) leads to an exponential decay in the rate of cell growth, resulting in longer cell cycle times proportional to *k*. Contact inhibition in our model is thus a consequence of physical constraint induced by tissue crowding (8), resulting in deviation of cell area from its preferred value in isolation (22).

The number of cells in the tissue, *N*, change in time due to the rules implemented in the decision-making layer. *N* can increase though cell divisions, which is regulated by a G1 sizer model (21, 22). In this model, a cell is in the G1 phase until its area is greater than a threshold value *A*_*S*_ (Fig. 1A-R1), at which point it transitions to the S/G2/M phase which lasts for a fixed time period τ_0_ (a timer). After a time τ_0_ in the S/G2/M phase, the cell divides into two cells (Fig. 1A-R2), upon which its target area is set to the current area. Both τ_0_ and *A*_*S*_ are drawn from a Gaussian distribution, with mean and standard deviation defined in Table 1.

**Table 1:** Default parameters used in model simulationsh.

| Parameter | Numerical value (units) | Description |
| --- | --- | --- |
| $k_B T$ | $1.95 \times 10^{-4} (\lambda A_S^2)$ | Thermal energy scale of Metropolis-Hastings algorithm |
| $J_{cc}$ | $1.17 \times 10^{-4} (\lambda A_S^{3/2})$ | Cell-cell contact energy |
| $J_{cm}$ | $1.96 \times 10^{-4} (\lambda A_S^{3/2})$ | Cell-medium contact energy |
| $\lambda$ | 1 | Cell area elastic modulus |
| $G$ | $1.16 (A_{S_0}/\tau_0)$ | Growth rate of an isolated cell |
| $k$ | $1.897 \times 10^5 (A_S^{-2})$ | Sensitivity to crowding |
| $\tau_0$ | $225 \pm 5.625$ (Monte Carlo Steps) | S/G2/M timer |
| $A_S$ | $1377 \pm 34.4$ (pixels) | G1 sizer threshold |
| $p_{apo,max}$ | 0.0015 | Maximum probability of apoptosis |
| $\rho_{1/2}$ | $1.5015 (A_S^{-1})$ | Cell density at half-maximum probability of apoptosis |
| $\alpha$ | $0.968 (A_S)$ | Sensitivity to density-dependent apoptosis |

On the other hand, the number of cells in the tissue can decrease via cellular apoptosis and extrusion. The model considers a density-dependent probability of apoptosis, *P*_apo_, which increases with the local cell density *ρ* (9, 37, 38) (Fig. 1A-R3)

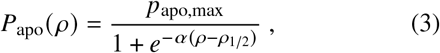

where *p*_apo,max_ is the maximum probability of apoptosis, *ρ*_1/2_ is the cell density at half-maximum probability and *α* is the sensitivity to density-dependent apoptosis. The parameters *p*_apo,max_, *ρ*_1/2_ and α are determined by fitting apoptosis probabilities calculated from cell competition experiments performed by Bove et al. (9). In our simulations, local cell density was defined as the sum of inverse cell areas within the immediate neighborhood of a given cell *α* (9, 26),

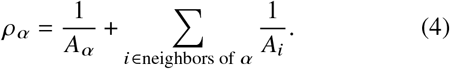

After entering apoptosis, the cell’s target area is set to zero, causing the cell to gradually decrease in area until being removed. Finally, a cell is eliminated via live cell extrusion if its area is less than a quarter of the mean cellular area, i.e., *A*_*a*_ *<* (*A*_*a*_)/4 (Fig. 1A-R4). Taken together, the rate of cell proliferation in the tissue is given by 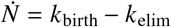, where *k*_birth_ corresponds to the rate of cellular divisions, and *k*_elim_ corresponds to the rate of cell elimination. Combining the physical and decision-making layers results in a predictive cell-based model (see Supplemental Information for details) that can simulate the spatiotemporal dynamics of a growing tissue in different biophysical contexts.

## RESULTS AND DISCUSSION

### Collective behaviors in growing epithelial colonies

To validate the predictive capacity of our model, we used it to recapitulate experimentally observed collective and single-cell behaviors in growing epithelial colonies (8, 22). We simulated the growth of a planar epithelial tissue, confined within a square box of area ∼ 1600*A*_*s*_ (Videos S1,S2). The simulations were initialized with a single cell of area below the G1 sizer threshold *A*_*s*_. As a first step, we performed two ensembles of simulations: (1) tissue growth without apoptosis (*P*_apo_ = 0, Video S1), and (2) tissue growth with apoptosis, occurring with a density-dependent probability *P*_apo_ as defined in Eq. (3) (Video S2). Here we considered the limit in which the scale of the elastic force, 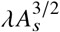, is larger than the tension between cells, *J*_*cc*_, and the tension between boundary cells and the external medium, *J*_*cm*_, with *J*_*cm*_ *> J*_*cc*_ (see Table 1 for a list of default parameter values). Consequently, the tissues grew while maintaining their initial isotropic state, with a roughly circular boundary, until they made contact with the simulation box (Fig. 2E-F, Videos S1,S2).

**Figure 2:**
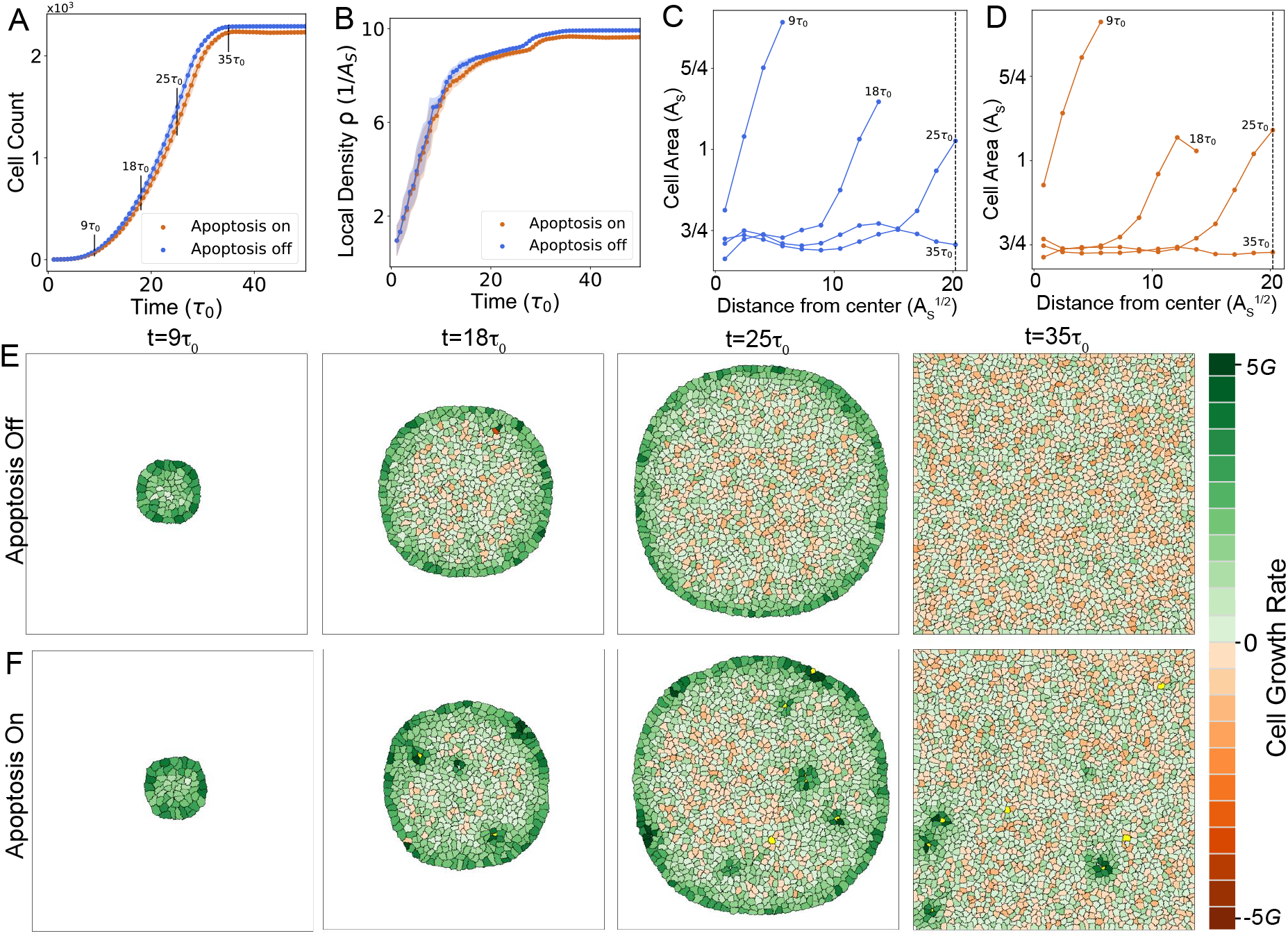
Characterization of collective behaviors in growing epithelial colonies. (A,B) Cell count vs time (A) and average cell density vs time (B) in colony growth simulations with apoptosis on (orange) and off (blue). The x-axis represents time measured in factors of τ_0_, the mean timer threshold. (C,D) Spatial distribution of single cell areas for colonies at different timepoints during growth, without (C) or with apoptosis (D). Each curve represents a colony’s spatially-averaged cell area as a function of distance from the center of the colony. The vertical dashed line is position of the boundary of the simulation box. (E,F) Time-lapse showing the morphologies of a growing epithelial colony without (E) or with (F) apoptosis. Cells are colored according to their growth rates. See Table 1 for a list of parameter values.

Our model accurately captured the experimentally measured trends in the dynamics of cell count and average cell density during epithelial colony growth and homeostasis (Fig. 2A-B) (9). During the growth phase of the tissue, constituent cells exhibited non-uniform sizes and growth rates, with a slowly growing bulk region surrounded by a rapidly proliferating periphery (Fig. 2E-F). Rapid replication of cells near the boundary have been previously observed in several experiments (8, 10, 13, 28). In our model this results from crowding-induced suppression of growth in the bulk of the tissue, which results in smaller cell size in the bulk compared to the periphery (Fig. 2C-D). With or without apoptosis, the simulated tissues grew until they reached confluence, filling the entire simulation box (Fig. 2E-F). This was followed by a state of homeostasis where the cell count and the average density did not change. In this state, the rates of cell division and elimination balanced to yield a net zero growth rate. Confluence led to uniform crowding throughout the tissue, resulting in the homogenization of the cellular areas and density (Fig. 2A-B).

In simulations without apoptosis, the rate of cell elimina-tion *k*_elim_ arises from live cell extrusions only. Homeostasis in these simulations is characterized by *k*_birth_ = *k*_elim_ = 0, where all the cells are arrested in the G1 phase, with *Aα < A*_*s*_, ∀*α*. By contrast, in simulations with apoptosis, the elimination rate is dependent on both live cell extrusions and apoptosis, the latter being a density-dependent probabilistic process. Therefore, homeostasis is characterized by a dynamic balance between birth and elimination, with *k*_birth_ = *k*_elim_ ≠ 0. Cell density-dependent apoptosis events induced cellular growth in neighborhood of a dying cell (Fig. 2F), leading to increased cell size, lower values of cell count and average cell density (Fig. 2A-B). Furthermore, cell elimination events increased the time required to reach homeostasis (Fig 2A), promoting continuous cell turnover and growth in the tissue.

### Single-cell dynamics underlying epithelial growth and homeostasis

Spatial regulation of cell proliferation is a direct consequence of how cell cycle and growth are regulated across a tissue (Fig. 2E-F, Fig S1). Here we characterize the single-cell growth and size dynamics underlying the growth of an epithelial colony. A major regulator of single-cell growth rate is local crowding by other cells. Growth rate of cells in the bulk of the tissue reduced over time due to an increase in local cell density, leading to a transition in the mitotic behavior of cells. During the early phase of growth (up to *t*≈ 7τ_0_), cells exhibited timer-like behavior with divisions occurring at regular intervals of *τ*_0_, due to the cell size being larger than the G1 sizer threshold *A*_*s*_ (Fig. 3A). During this period, cell size distribution was unimodal (Fig. 3C-D), with a mean value *Ā > A*_*s*_. Mean cell area, *Ā*, decreased in time since the amount of growth accrued in the timer (S/G2/M) phase *G*τ_0_ ∼ 1.16*A*_*s*_ *< Ā*, was less than the amount of cell size reduction via division. This phenomenon of size-reductive divisions recapitulates experimental observations in growing MDCK colonies (8, 22).

**Figure 3:**
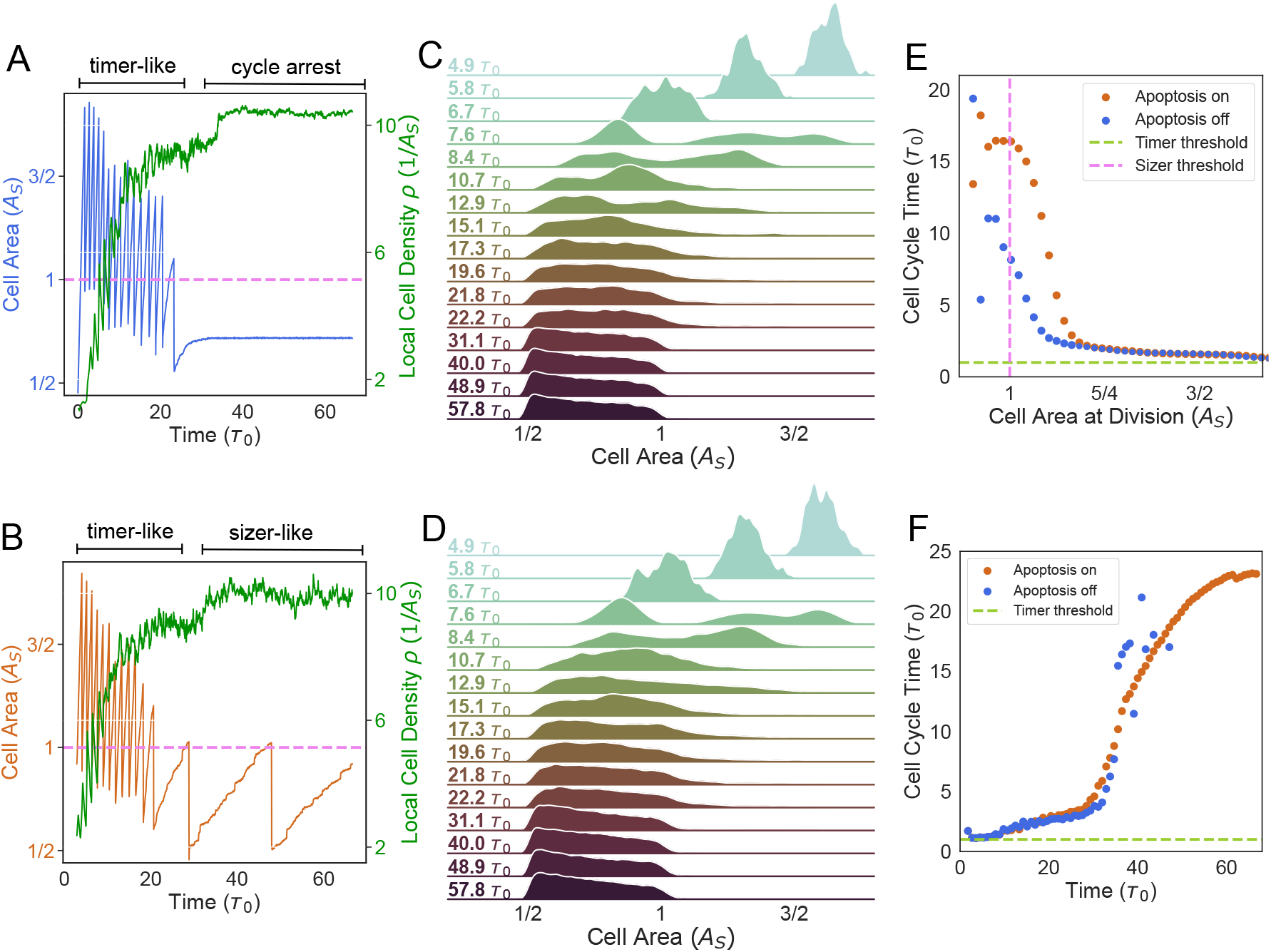
Single-cell growth and cell cycle dynamics in growing tissues. (A,B) Single-cell area trajectories (A - blue trajectory in simulations without apoptosis, B - orange trajectory in simulations with apoptosis) and average local cell density vs time (green). In (A), cell cycle exhibited timer-like behavior until crowding led to cell cycle arrest. In (B), cells initially exhibited timer-like behavior, and then transitioned to a sizer-like control of cell cycle. (C,D) Evolution of cell area distribution in simulations without apoptosis (C), and in simulations with apoptosis (D). (E) The average cell cycle time as a function of mean cell area at mitosis. The dashed violet vertical line is the mean sizer threshold and the dashed green horizontal line is the mean timer threshold. (F) The average cell cycle time as a function of time. Division events ceased soon after confluence for simulations without apoptosis (blue). Simulations with apoptosis (orange) showed a linear increase in cycle time until plateauing near the end.

Once cell size in the population fell below *A*_*s*_ (Fig. 3C-D, *t >* 7*τ*_0_), two sub-population of cells emerged: one with mean size above *A*_*s*_ that would divide after time *τ*_0_ in the G2 phase, and another with mean size below *A*_*s*_ that would be growing in the G1 phase, and therefore would divide in a time longer than *τ*_0_. This behavior is evident in the cell area distribution, that changed from a unimodal to a bimodal shape (see *t*∼7.6*τ*_0_ in Fig. 3C-D). However, as cell crowding increased via an increase in the local cell density (Fig. 3A-B), cell area decreased over several generations, narrowing the area distribution (Fig. 3C-D). A consequence of crowding-induced slowdown of single cell growth is a negative correlation between cell area and cell cycle time (Fig. 3E-F), as observed experimentally (8).

Post-confluence (*t >* 35 *τ*_0_), cell growth was either non-existent or slow due to the lack of available free space. In simulations without apoptosis, cells underwent complete cycle arrest (Fig. 3A), as a result of the lack of tissue turnover to free up space. By contrast, in simulations with apoptosis, the rates of growth and division were limited by the available free space which appears after cell elimination. Consequently, the cell cycles progressed at a very slow rate (*t* ≥ 20τ_0_ in Fig. 3B), primarily determined by the rate of apoptosis within the tissue. These slowly growing cells were stuck in the G1 phase for the majority of the cell cycle and divided relatively fast (fraction of the cycle spent in G1 vs S/G2/M) after exceeding the G1 size threshold *A*_*s*_, thereby exhibiting sizer-like behavior.

### G1 sizer sets homeostatic cell density and turnover rate

Having validated our model predictions for collective and single cell behaviors against experimental observations, we turned to assessing the roles of cell-scale parameters on the emergent tissue-level properties during growth and homeostasis. To this end, we varied the model parameters regulating cell growth and division dynamics, including the growth rate *G*, area elastic modulus *λ*, cell-cell adhesion energy *J*_cc_, sensitivity to crowding *k* and the G1 sizer threshold *A*_*S*_. As a first step, we studied the effect of these parameters on homeostatic tissue properties, including homeostatic cell density and turnover rate.

Prior studies have demonstrated that epithelial tissues possess a specific density they tend to restore after perturbations, indicating their tendency to maintain a preferred homeostatic density (14, 39, 40). However, the mechanisms underlying the regulation of this homeostatic density, which plays a pivotal role in maintaining tissue homeostasis (14, 38, 41) and cell competition (26, 37), remain poorly understood. Our simulations revealed that the G1 sizer threshold *A*_*S*_ is the only parameter that determines the homeostatic density (Fig. 4A,C). This can be intuitively understood from the fact that *A*_*s*_ regulates cell size limit in homeostasis (Fig. 3C-D) and cell density is related to the inverse of cell areas (Eq. 4). Cells larger than *A*_*S*_ would divide after the timer phase, regardless of other factors. Consequently, the rate of tissue turnover in homeostasis, which is set by the density-dependent rate of apoptosis (Eq. 3), is also set by the cell size threshold *A*_*S*_ (Fig. 4A). In particular, cells with a larger *A*_*S*_ assembled tissues with a slower turnover rate and a slower division rate compared to those with a smaller value of *A*_*S*_ (Fig. 4D). Therefore, cell size regulation plays a major role in setting the physical properties of tissues at homeostasis, including cell density and turnover rate. By contrast, other physical parameters (such as growth rate, elasticity, contact energy and crowding) do not regulate homeostatic density and turnover rates.

**Figure 4:**
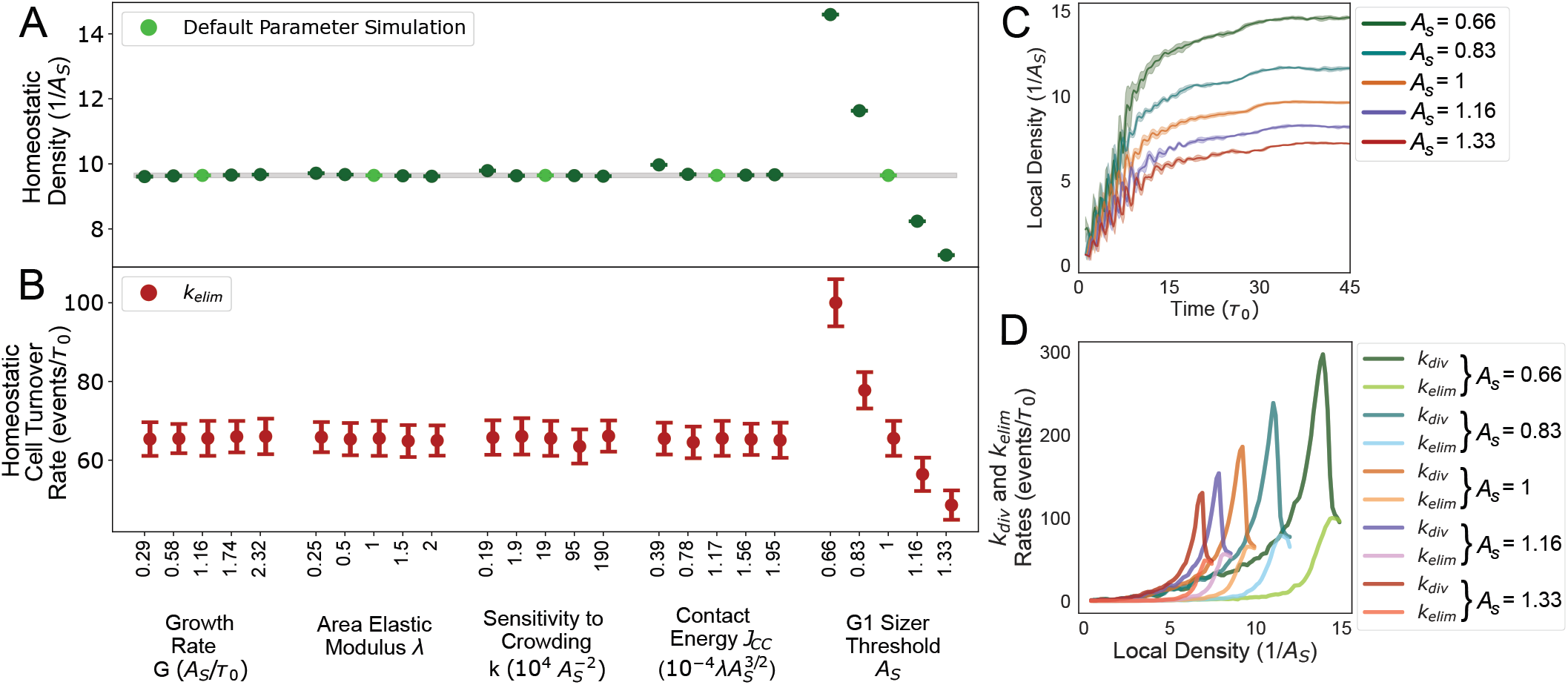
G1 sizer regulates homeostatic density and tissue turnover rate. (A) Average cell density at homeostasis as a function of various simulation parameters. The G1 sizer threshold, *A*_*S*_, sets the homeostatic density of the tissue. The default parameter values are identified with light green solid circles. (B) Average rate of tissue turnover at homeostasis, given by the elimination rate *k*_elim_, as a function of various simulation parameters. (C) The average local cell density as a function of time for various values of the sizer threshold *AS*. (D) Average rates of cell division and elimination as a function of local cell density for various values of the sizer threshold *A*_*S*_.

### Cellular pressure regulates tissue growth rate

Having analyzed tissue properties in the homeostatic state, we proceeded to examine how model parameters influenced tissue growth dynamics (Fig. 5A). For this purpose, we systematically varied cell mechanical parameters (*λ, k* and *J*_cc_) and the factors controlling cell size and growth (*G* and *A*_*S*_), analyzing their impact on tissue growth rate. To quantify the timescale of tissue growth *t*_growth_, we measured the time taken by the cell colony to reach the size of the confinement. We found that the timescale *t*_growth_ was primarily influenced by the cellular growth rate *G*, compressibility *λ*, and the sensitivity to crowding *k* (Fig. 5B). In contrast, variations in cell-cell contact energy *J*_cc_ and cell size threshold *A*_*S*_ had minimal impact on *t*_growth_. As a result, the rate of tissue growth, which is inversely proportional to *t*_growth_, is predominantly determined by cell mechanical parameters. Moreover, we observed that the regulation of tissue growth rate is independent of homeostatic density (regulated by *A*_*S*_), contrary to assumptions made by other models (41).

**Figure 5:**
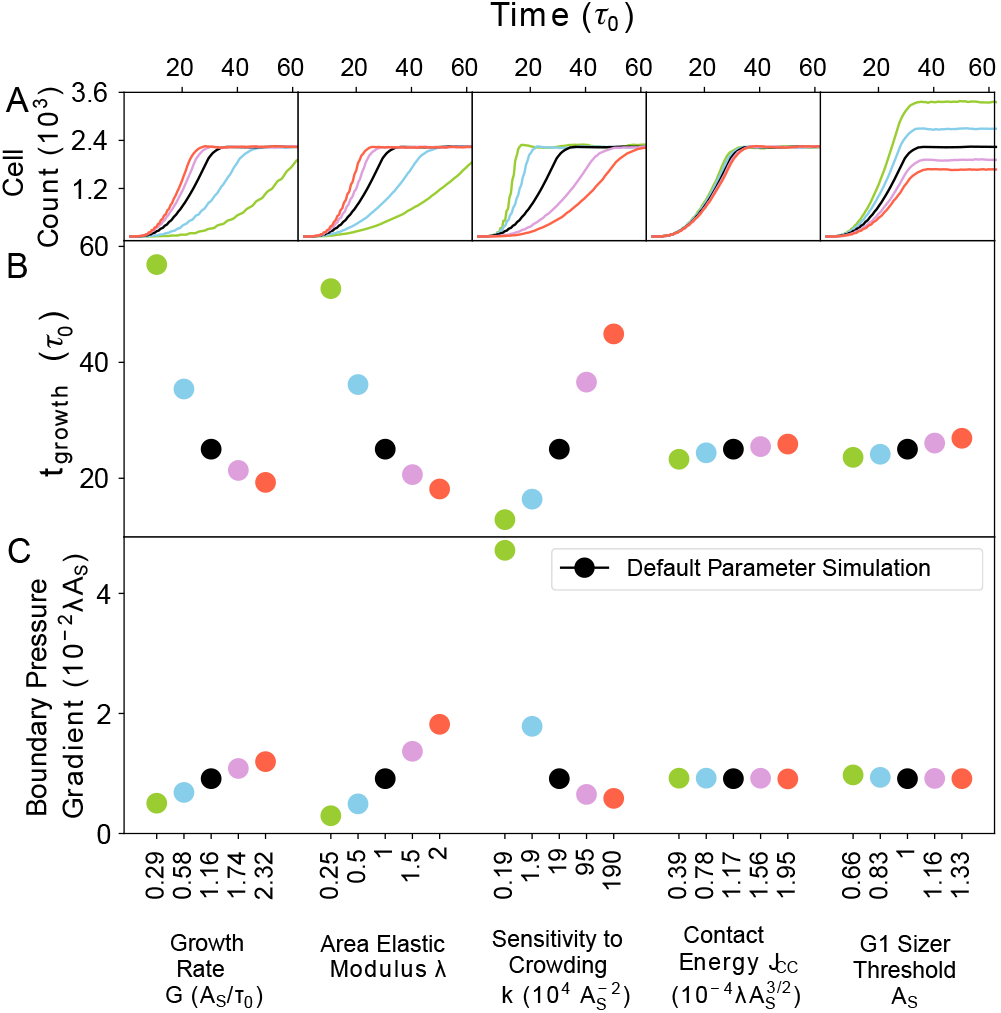
Sensitivity to crowding determines the rate of tissue growth, independent of homeostatic cell size. We quantify the timescale of tissue growth (*t*_growth_) as the time taken by the cell colony to reach the size of the confinement. (A) Cell count vs time (in units of *τ*_0_) for different values of the single-cell parameters {*G, λ, k, J*_cc_, *A*_*S*_}. (B) The timescale of tissue growth, *t*_growth_ (in units of *τ*_0_), for different values of the parameters {*G, λ, k, J*_cc_, *A*_*S*_}. Increasing *G* and *λ* results in a decrease in *t*_growth_, while *t*_growth_ increases with the sensitivity to crowding (*k*). By contrast, 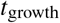 is weakly dependent on *J*_cc_ or *A*_*S*_. (C) Pressure gradient at the boundary of the cell colony vs single-cell parameters {*G, λ, k, J*_cc_, *A*_*S*_ }. The correlation between pressure gradient and 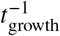 reveals that cellular pressure regulates the rate of colony growth.

The mechanical parameters that govern *t*_growth_ directly influence the cellular response to crowding by regulating cell pressure (Fig S2). The cell pressure *p*_*α*_ is given by the equation:

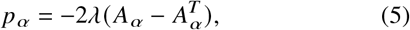

where A α is the cell area and 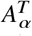 denotes the target cell area. The pressure gradient between the interior and exterior of the colony boundary exerts radial outward forces on the cells (Fig S2), with this pressure gradient predominantly controlled by the cell mechanical parameters *G, λ*, and *k* (Fig. 5C). The magnitude of the pressure gradient at the tissue boundary characterizes the tissue’s sensitivity to crowding, consequently impacting the net growth rate (Fig. 5C).

To conceptually understand the roles of cell-level (micro-scale) interactions on tissue-level (macro-scale) growth dynamics, we developed a simple mathematical model of a growing elastic tissue. Here we neglected spatial variations in proliferation and modeled the entire tissue as a single continuous medium of area *A*, preferred area *A*_0_, and stress relaxation rate *λ*. The stress relaxation rate is related to the elastic modulus governing cellular resistance to area changes. Then, the colony size evolves as

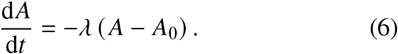

Here *A*_0_ can be interpreted as the total area of the cells if they grew in isolation, such that stress relaxation occurred instantaneously. For isolated cells, cellular duplication happens at a constant rate defined as *G a*_0_, where *G* is single-cell growth rate and *a*_0_ is the area of a newborn cell proliferating in isolation. Instead, if cells are not isolated, the proliferation rate is not constant, and decreases as a function of the absolute value of pressure, as in the microscopic model. In this case divisions occur at a rate (*G*/ *a*_0_) exp [*k A*−*A*_0_)^2^], with *k* a sensitivity to crowding. In order to model a tissue cells growing in a confinement, the equation governing the preferred colony area *A*_0_ is given by

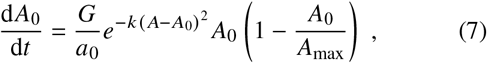

where *A*_max_ is the area of the confinement. Therefore, in the homeostatic steady-state, colony area will be *A* = *A*_0_ = *A*_max_. We solved Eqs. (6) and (7) numerically, starting with an initial condition of *A*(*t* = 0) = *A*_0_(*t* = 0) = *a*_0_, for different values of the growth rate *G*, stress relaxation rate *λ*, and crowding sensitivity *k* (Fig. 6A). In the continuum model simulations, we scaled area units by *a*_0_ and time units by 1/ *λ*, with *A*_max_ set to 800*a*_0_. We defined *t*_growth_ as the time it takes for the colony area to reach 0.95*A*_max_. The continuum model solutions closely replicated the trends observed in the cellular Potts model simulations (Fig. 5) for both the dynamics of colony area (Fig. 6A) and *t*_growth_ (Fig. 6B) in relation to *G, λ*, and *k*. Particularly, we found that *t*_growth_ increased with decreasing tissue pressure, induced by an increase in *k*, or a decrease in *G* or *λ* (Fig. 6B). This showed that mechanical pressure within the tissue served as the primary regulatory factor for its growth rate.

**Figure 6:**
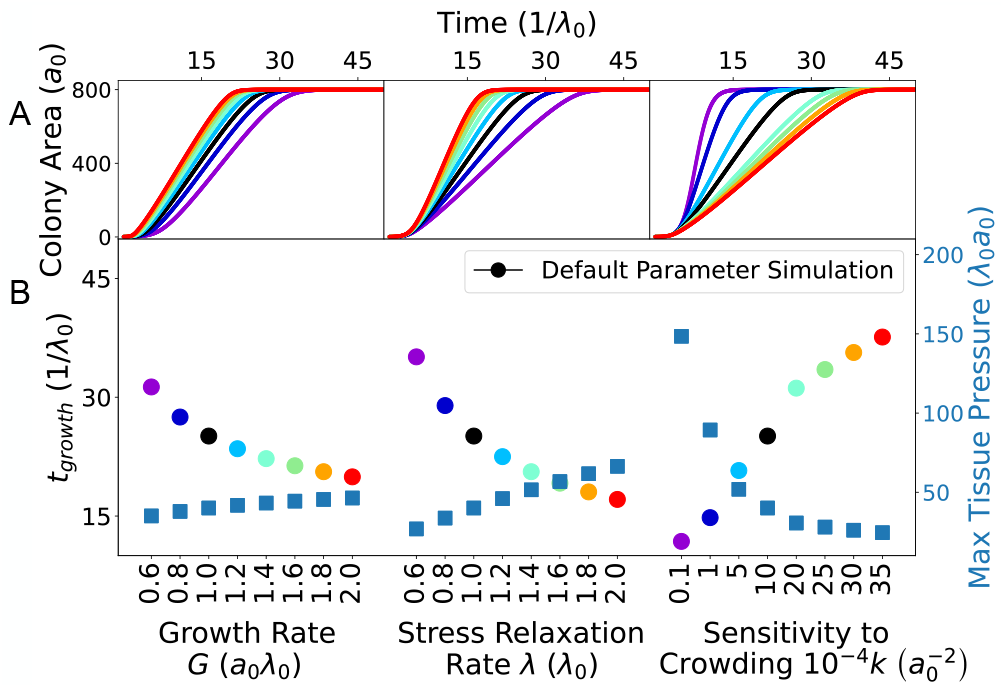
Predictions of a continuum model for a growing cell colony. Model results for the (A) colony area *A*_*c*_ *t*, (B) timescale of colony growth *t*_growth_, and (B) maximum tissue pressure, for varying growth rate *G*, stress relaxation rate *λ*, and crowding sensitivity *k*. The timescale of colony growth is defined as *A*_*c*_ (*t*_growth_) = 0.95*A*_max_, where *A*_max_ = 800*a*_0_ is the maximum area occupied by the colony, given by the area of the confinement.

### Cellular elasticity and sensitivity to crowding control the emergent patterns of cell proliferation

Crowding-induced suppression of growth results in an uneven pressure distribution across the tissue, with high pressure in the center and low pressure at the tissue boundary (Fig S2). As a consequence of this pressure gradient, tissue growth becomes nonuniform, with a bulk region experiencing a low proliferation rate (proportional to the tissue turnover rate) compared to the external region where proliferation occurs at a higher rate (Fig. 2). However, experimental observations in both *in vivo* and *in vitro* growing epithelial tissues have revealed diverse growth patterns, including scenarios where the bulk region of the tissue exhibits dynamic growth at the cellular level (13, 22, 28, 29). Therefore, we further investigated how our model could reproduce various tissue-level growth patterns by modulating cellular pressure distribution, which is regulated by the compressibility *λ* and the sensitivity to crowding *k*.

First, we conceptually explain the role of pressure on the spatial distribution of growth. Consider a growing cell colony that originates from a single isolated cell. Initially, this cell is surrounded only by the medium, which doesn’t hinder its growth, allowing it to rapidly expand at a rate *G*. As the cell divides and gives rise to two new cells, each newborn cell finds itself surrounded by two different environments - neighboring cells and the medium. Due to the similarity in sensation between these environments, the growth rate remains similar for both types of cells. As multiple divisions occur, the tissue becomes a mix of two cell types: bulk cells entirely surrounded by other cells and boundary cells in contact with the external medium. If cell growth is sensitive enough to the pressure exerted by neighboring cells, controlled by *k* and *λ* (Eq. 5), the growth rate of these two cell groups will significantly differ. Boundary cells, experiencing no external pressure at the tissue boundary, will proliferate faster than bulk cells due to the lesser pressure from the surrounding cells.

To investigate the impact of cell pressure on tissue growth patterns, we conducted simulations of growing tissues, varying elasticity *λ* and contact inhibition *k* (Videos S3,S4,S5). Specifically, we analyzed the distribution of cell growth rate and cell cycle phase just before the colony reached the boundary of the simulation box (Fig. 7). For higher values of *λ* and *k*, we observed a strong radial gradient of growth rate, with a bulk region largely arrested in the G1 phase (Video S3). Reducing *λ* reduced the pressure difference at the tissue boundary, resulting in slower growth in the outermost ring and inducing an inward pushing force. This inward force caused contraction of bulk cells (negative area growth rates in Fig. 7). Consequently, cell cycles were extended throughout the tissue, with most cells predominantly in the G1 phase (cyan color in Fig. 7). Conversely, decreasing *k* enhanced the growth rate of bulk cells while homogenizing growth rates across the tissue (Video S4). For very low values of *k*, uniform growth was observed throughout the tissue. Remarkably, in this scenario, all cells continued to grow as if they were not constrained, leading to a substantial fraction of cells in the timer phase (magenta color in Fig. 7, Fig S3). For intermediate values of *k* and *λ*, we observed spatially mixed patterns of proliferation, with alternating rings of G1 and G2 phases often present (Video S5). Overall, the fraction of cells growth arrested in G1 phase increased with increasing *k* and decreased with *λ* (Fig S3). Hence, cell mechanical parameters, such as elasticity and sensitivity to crowding, significantly influence tissue-scale cell proliferation patterns by manipulating pressure within the tissue.

**Figure 7:**
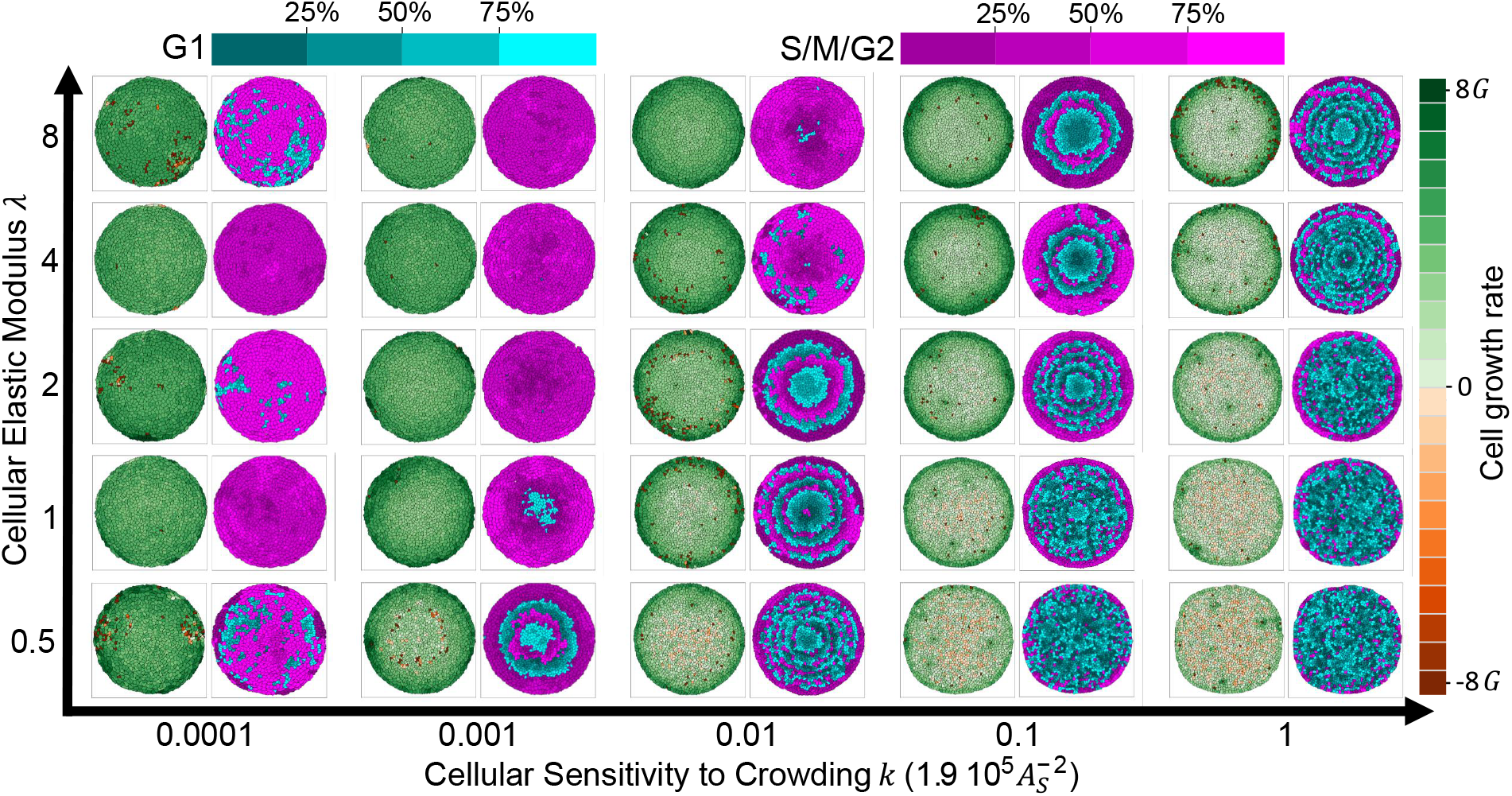
Spatial patterns of tissue growth depicted for different values of cellular elastic modulus (*λ*) and sensitivity to crowding (*k*). Each snapshot is captured just before contact with the boundary occurs, allowing an insight into patterns driven by internal tissue dynamics rather than the physical effects of confinement. The snapshots are color-coded to represent growth rate (left, green and red) and cell cycle phase (right, cyan and fuchsia). As the elastic modulus (*λ*) is varied from high to low or the sensitivity to crowding (*k*) changes from low to high, three distinct growth patterns emerge: uniform growth (where the majority of the tissue is in the same cell cycle phase), locally uniform growth (resulting in ring patterns due to cells having similar phases as their neighbors), and boundary growth (where only the outer ring of cells is synchronized in phase, while inner cells stochastically grow at the tissue turnover rate).

## CONCLUSIONS

Our current understanding of the mechanisms regulating single-cell growth and division is extensive (15, 16). However, the coordination of these processes across different spatiotemporal scales, which is crucial for the regulation of tissue proliferation, remains poorly elucidated.

To address this, we developed a multi-layered modeling framework using the Cellular Potts Model (31, 34). This model integrates single-cell decisions governing cell fate with their mechanical interactions within the tissue. We incorporated rules for cell growth, cell cycle regulation, contact inhibition of proliferation, and cell elimination via cell-density-dependent apoptosis based on existing knowledge of cell-level proliferation rules (8, 9, 21, 22, 26). Specifically, the cell cycle consists of an initial G1 sizer phase followed by an S/M/G2 timer phase. Our model successfully reproduced various experimentally observed single-cell behaviors, including the transition of mitotic behavior from *timer-like* to *sizer-like* control of cell size, regulated by crowding. Moreover, the model captured emergent characteristics seen in previous experiments, such as tissue homeostasis (9, 41), the cell-size-dependent cycle time in confluent mammalian cells (8, 21, 22), the tissue crowding-induced narrowing of cell area distributions and increased cell cycle time (8), as well as diverse spatial patterns of tissue growth (22, 28–30).

Through *in silico* experiments, we have identified the G1 size threshold, *A*_*S*_, as the critical parameter determining the homeostatic cell density. This size threshold governs the rate of tissue turnover in homeostasis, as apoptosis rates depend on local cell density. The specific molecular mechanisms encoding the G1 size threshold value remain unclear, although a recent study indicated a link between the size threshold and genome size (22). Consequently, perturbations to genome size in epithelial tissues could serve to test our predictions regarding the role of cell size in regulating homeostatic cell density and tissue turnover.

Furthermore, our simulations demonstrated that cellular attributes like elasticity, growth rate, and sensitivity to crowding did not directly impact homeostatic density and turnover rate. However, these parameters did influence the dynamics of tissue growth before reaching homeostasis. Particularly, tissue-level growth rate was positively controlled by cell-level growth rate and elasticity, while being negatively influenced by the sensitivity to crowding. Using a continuum model, we established that this control is achieved by adjusting the pressure within the growing tissue. Significantly, we discovered that the interplay between cellular stiffness and sensitivity to crowding, with opposing effects on tissue pressure, played a crucial role in determining spatial growth patterns. This interplay resulted in the emergence of diverse spatial patterns of growth, ranging from uniform growth to localized growth. These findings offer insights into potential pressure-related mechanisms that could be harnessed to regulate tissue growth. One intriguing scenario involves considering a buffer tissue surrounding the growing tissue. Our work provides a guide for identifying which cell-level mechanisms the buffer tissue should modify to regulate the invasion of the inner colony.

## AUTHOR CONTRIBUTIONS

S.B. designed and supervised the study. S.B. and L.C.C. developed the computational model. L.C.C carried out simulations and analyzed the data. F.P. developed the continuum theory. L.C.C., F.P., and S.B. wrote the article.

## ACKNOWLEDGEMENTS

We thank Daniel Gradeci for useful discussions. SB acknowledges support from the National Institutes of Health (NIH R35 GM143042).

