## Supplemental Information for "Mechanical control of cell proliferation patterns in growing tissues"

### SUPPLEMENTAL METHODS

#### Cellular Potts Model

Our cell-based model for a growing tissue integrates two computational components: (1) a physical layer that simulates cell mechanics with the Cellular Potts Model (CPM) [1], and (2) a decision-making layer that defines rules for cell growth, division, and apoptosis. The CPM is a versatile computational framework for simulating biological systems such as cell growth, migration, and tissue formation. The CPM runs a Monte Carlo simulation with the Metropolis-Hastings algorithm [2, 3] to minimize the free energy of the system. A common choice of the Hamiltonian [1] includes terms that represent contact energies and an area elastic energy, as in

$$E = \sum_{\substack{i,j \\ \text{neighbors}}} J_{ij}\left(\tau_i, \tau_j\right) + \lambda \sum_{\alpha \in \text{cells}} \left(A_\alpha - A_\alpha^T\right)^2 , \tag{S.1}$$

The contact energy, $J_{i,j}$, is given by the difference between the cellular surface tension and intercellular adhesion energy. This energy is represented in the Hamiltonian by a component that is proportional to the number of pixels shared between cells or between a cell and the medium. The sum goes over all adjacent lattice sites $i$ and $j$, ignoring self-adhesion. The inner term, $\tau$, gives the cell type at the lattice site $i$ and $j$ respectively. A value for contact energy is defined between each cell type, which for our simulations consists of interactions between cells and medium $J_{\text{cell-cell}} = J_{\text{cc}}$, $J_{\text{cell-medium}} = J_{\text{cm}}$, and $J_{\text{medium-medium}} = J_{\text{mm}}$. The second term in the Hamiltonian is the area elastic energy. This is given by the product of the area elastic modulus, $\lambda$, with the sum of squared differences between the actual area of the cells, $A_\alpha$, and their target areas, $A_\alpha^T$. These two mechanical energy components regulate individual cell shape and size, in the absence of other rules. The CPM simulations were run using CompuCell3D [4], a software package that handles all of the computations necessary for the CPM and the Metropolis-Hastings algorithm.

#### Rules for active cell behaviors

The decision-making layer of the model encodes probabilistic rules for active cell behaviors including changes in cell target area, cell cycle regulation, and cell elimination.

*Dynamics of cell growth* – Cell growth dynamics are implemented by increasing the cell's target area as

$$\frac{\mathrm{d}A_\alpha^T}{\mathrm{d}t} = Ge^{-k(A_\alpha - A_\alpha^T)^2}, \tag{S.2}$$

where $G$ is the growth rate in the absence of crowding, $k$ quantifies cellular sensitivity to crowding, and $A_\alpha$ is the current area of cell $\alpha$. In the presence of crowding, a cell's area will deviate from its target area, and this deviation is directly proportional to the strength of contact inhibition. Consequently, cellular growth exhibits an exponential decline as crowding intensifies, ultimately leading the tissue to enter an arrested state.

*Cell cycle regulation* – We implemented a two-phase cell cycle model, comprising a G1 growth phase operating through a sizer mechanism, followed by an S/G2/M timer phase. When a cell is born, it is assigned a G1 area threshold $A_S$ and an S/G2/M phase timer $\tau$. The cell remains in the G1 phase until its area exceeds $A_S$. Afterward, it enters the S/G2/M phase, which lasts for a duration of $\tau$. Upon completion of the S/G2/M phase, the cell immediately undergoes division, perpendicular to the semi-major axis, resulting in two daughter cells, both of which enter the G1 phase.

*Mechanisms for cell elimination* – TIn the simulation, cell elimination involves two mechanisms: programmed cell death (apoptosis) and live-cell extrusion. Experimental data have shown that increasing cell density leads to a higher apoptosis rate [5–7]. We defined the local cellular density as the sum of inverse cell areas within a local neighborhood of a cell. The density of cell $\alpha$ is given by:

$$\rho_\alpha = \frac{1}{A_\alpha} + \sum_{i \in \text{neighbors of } \alpha} \frac{1}{A_i}. \tag{S.3}$$

To model the probability of apoptosis in wild-type MDCK cells as a function of local cell density [7], we fit the data to a sigmoid curve:

$$P_{\text{apo}}(\rho) = \frac{p_{\text{apo,max}}}{1 + e^{-\alpha(\rho - \rho_{1/2})}}. \tag{S.4}$$

In the simulations, the probability of apoptosis is calculated every 10 Monte Carlo steps, which, on the time scale of the timer threshold, reproduces the experimentally observed probabilities. When a cell is eliminated through apoptosis, its target area is set to zero, causing it to shrink rapidly. During this process, surrounding cells grow to occupy the space left by the apoptotic cell.

In addition to apoptosis, cells can also be eliminated via live cell extrusions. This occurs when a cell's area significantly decreases compared to the average population area. In experiments [5], extrusion happens when tissue experiences compressive stress, causing a cell to delaminate from the substrate and eject from the monolayer. In our simulations, a cell whose area falls below $\langle A \rangle / 4$ (where $\langle A \rangle$ is the average area of the colony) is immediately removed to capture the short time scale over which extrusion occurs [5, 8].

### Choice of Default Model Parameters

Converting lattice units in the CPM framework to physical units is essential for comparing our simulations with experimental observations. There are two key conversions: translating pixels and Monte Carlo Steps (mcs) into SI units for distance and time. To achieve this, we compared the experimentally reported values for average cell volume during subconfluence for MDCK cells [9–11] with the value for average subconfluent colony area in our simulations, $\bar{A}_{SC} = 3200$ pixels. This comparison yielded the conversion factor: 1 pixel $= (0.13 \pm 0.03)\mu m^2$. Applying this factor to the G1/S size threshold, we obtained $A_S = 1377$ pixels $= (179.01 \pm 41.31)\mu m^2$. We normalized simulation time with the average cycle time of an uncrowded cell, i.e., the cell's timer duration $\tau_0$. Experimental data for typical epithelial cell types suggest that the time for the S/G2/M phase is approximately $t \approx 10$ hours [9, 11]. Keeping the timer threshold fixed at $\tau_0 = 225$ mcs provides the timescale for Monte Carlo steps.

The single-cell growth rate $G$ is sampled from a Gaussian distribution to determine the change in the target area at each Monte Carlo step (Eq. (S.2)). The mean value of $G$ is set such that isolated cells maintain the average subconfluent area $\bar{A}_{SC}$, $G_0 = \frac{1}{2}\frac{\bar{A}_{SC}}{\tau_0}$, and the standard deviation is set to 0.05 $G_0$. While $G$ sets cellular growth rate ($\mathrm{d}A^T/\mathrm{d}t$) in an isolated state, tissue crowding reduces the available space for growth to bulk cells, causing their actual area to deviate from the target area. This results in the domination of the exponential term of Eq. (S.2), $e^{-k(A-A^T)^2}$, and drives the growth $\mathrm{d}A^T/\mathrm{d}t$ to zero until the cell's area increases. The sensitivity to crowding, represented by the heuristic parameter $k$, quantifies the allowable deviation between a cell's area and its target area The default value, $k = 0.1$ pixels$^{-2}$, was determined empirically to ensure the tissue growth rate captures the expected growth dynamics over several cell cycles.

In our implementation of the CPM, there are four parameters within the Hamiltonian that set the energy scale; elastic energy has $\lambda$ and contact energy has $J_{\mathrm{cc}}$, $J_{\mathrm{cm}}$, and $J_{\mathrm{mm}}$. The parameter $\lambda$ represents the area elastic modulus for the cell, and we choose a default value $\lambda = 1$. The parameter $J$ represents the disparity between cell surface tension and intercellular adhesion, implying that higher $J$ values indicate lower cell-cell adhesion. Consequently, we chose $J_{\mathrm{cc}} = 6$ and $J_{\mathrm{cm}} = J_{\mathrm{mm}} = 10$, which allowed cells to adhere more strongly to each other than the medium, resulting in a circular morphology for the growing colony.

### Video Legends

**Video S1**. Simulation of a growing epithelial colony with apoptosis turned off. The simulation uses default parameters as listed in Table 1. Individual cells are color coded by their growth rates.

**Video S2**. Simulation of a growing epithelial colony with apoptosis turned on. The simulation uses default parameters as listed in Table 1. Individual cells are color coded by their growth rates.

**Video S3**. Simulation of a growing epithelial colony exhibiting boundary growth due to strong contact inhibition. The simulation uses default parameters as listed in Table 1, except elastic modulus $\lambda = 1$ and sensitivity to crowding $k = 1$. Left: Cells are color coded by their growth rates, Right: Cells are color-coded by the cell cycle phase (blue: G1 phase, fuchsia: S/G2/M phase).

**Video S4**. Simulation of a growing epithelial colony exhibiting uniform bulk growth due to weak contact inhibition. The simulation uses default parameters as listed in Table 1, except elastic modulus $\lambda = 4$ and sensitivity to crowding $k = 0.0001$. Left: Cells are color coded by their growth rates, Right: Cells are color-coded by the cell cycle phase (blue: G1 phase, fuchsia: S/G2/M phase).

**Video S5**. Simulation of a growing epithelial colony exhibiting patterned growth at intermediate contact inhibition. The simulation uses default parameters as listed in Table 1, except elastic modulus $\lambda = 2$ and sensitivity to crowding $k = 0.01$. Left: Cells are color coded by their growth rates, Right: Cells are color-coded by the cell cycle phase (blue: G1 phase, fuchsia: S/G2/M phase).

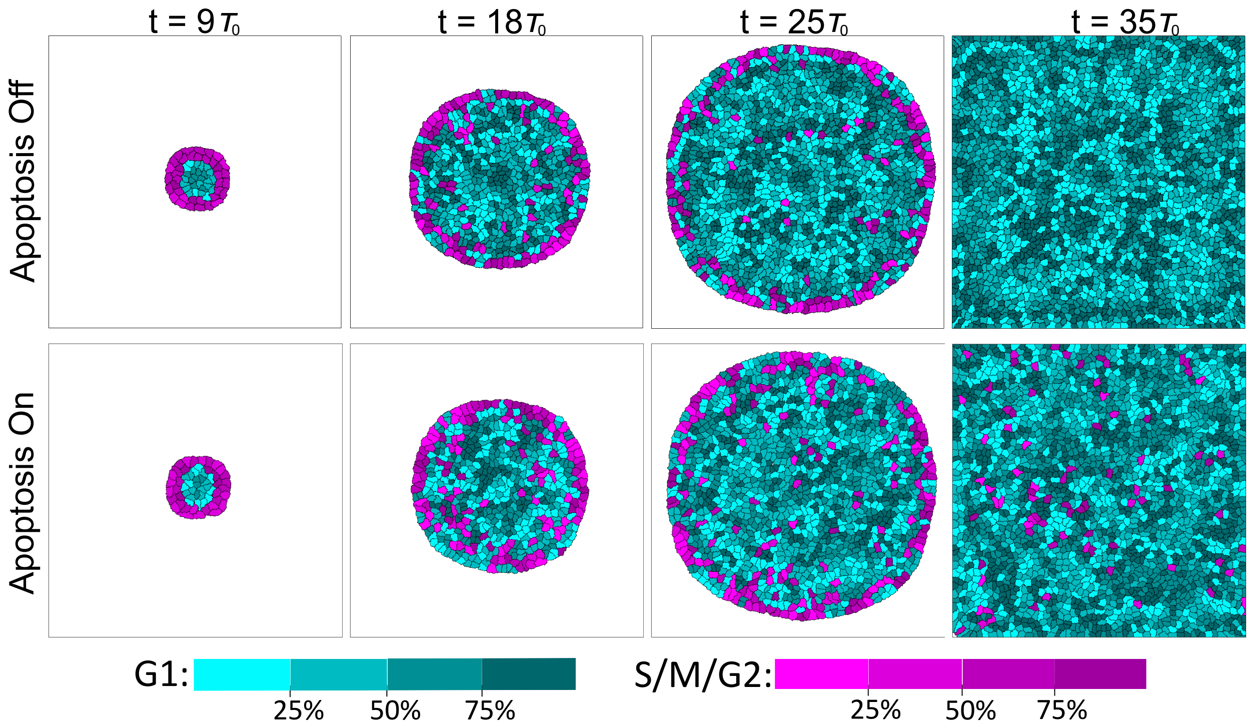


**Figure S** 1. Time-lapse simulation outputs of a growing colony, color-coded by the cell cycle phase (blue: G1 phase, fuchsia: S/G2/M phase). In both simulations, less crowded outer cells grew more rapidly, enabling them to transition from G1 to S/G2/M phases. By $t = 25\tau_0$, we observed limited, yet noticeable, growth within the bulk of the tissue, with the simulation using apoptosis showing more growth. At $t = 35\tau_0$, the tissue without apoptosis remained in the G1 phase, while the tissue with apoptosis exhibited turnover as dying cells created space for growth and cell cycle progression.

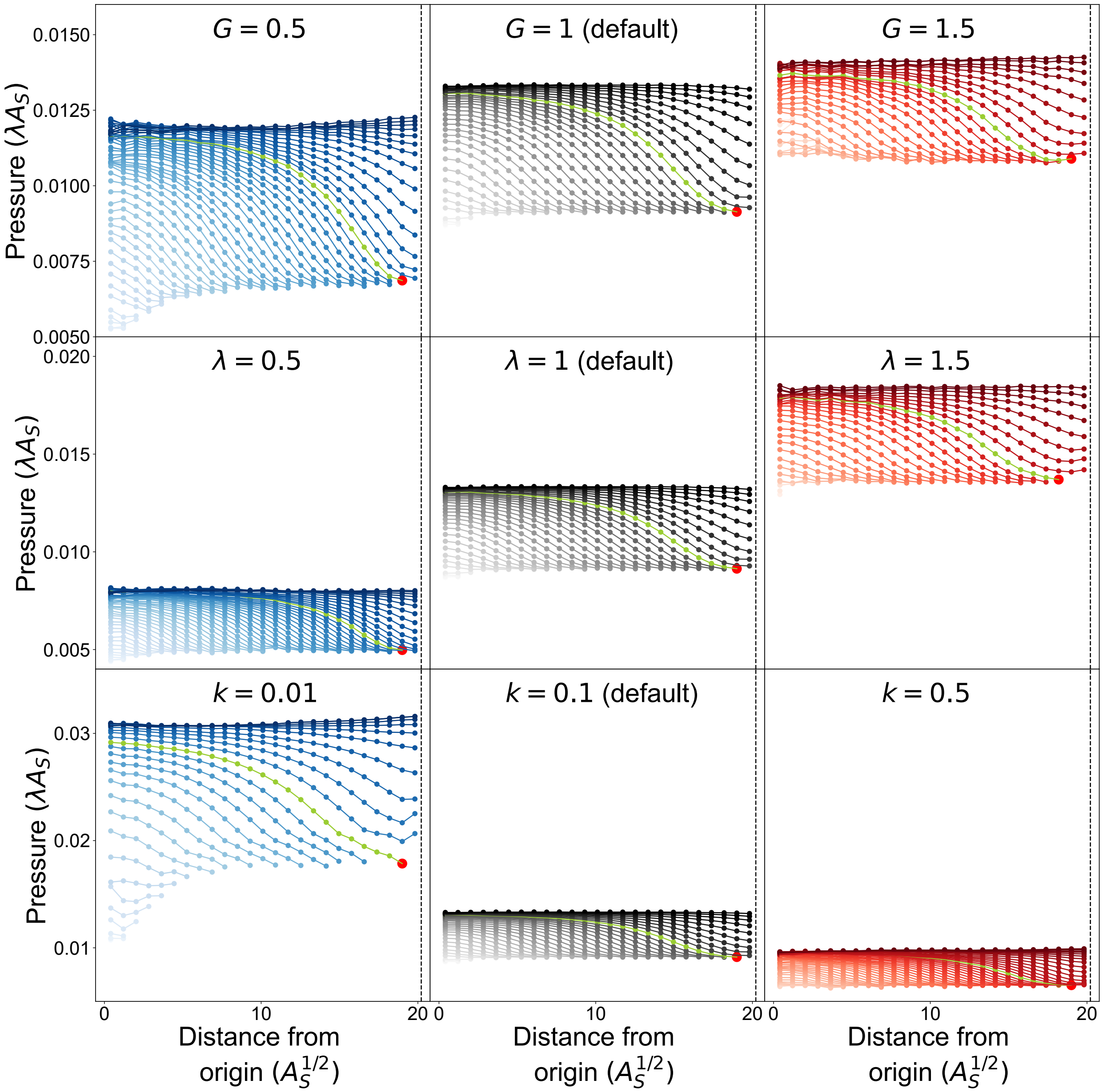


**Figure S** 2. Spatial distribution of cell pressure at various timepoints during colony growth, for different values of growth rate ($G$), sensitivity to crowding ($k$), and elasticity ($\lambda$). Each curve, separated in time by $8/9\tau_0$, depicts the radially-averaged cell pressure as a function of distance from the colony's center. The vertical dashed line indicates the position of the colony edge. The green curve shows the pressure distribution just before the tissue contacts the nearest boundary. The red point at the end of the green curve corresponds to the value of the boundary pressure gradient plotted in Fig 5 for each set of these parameters.

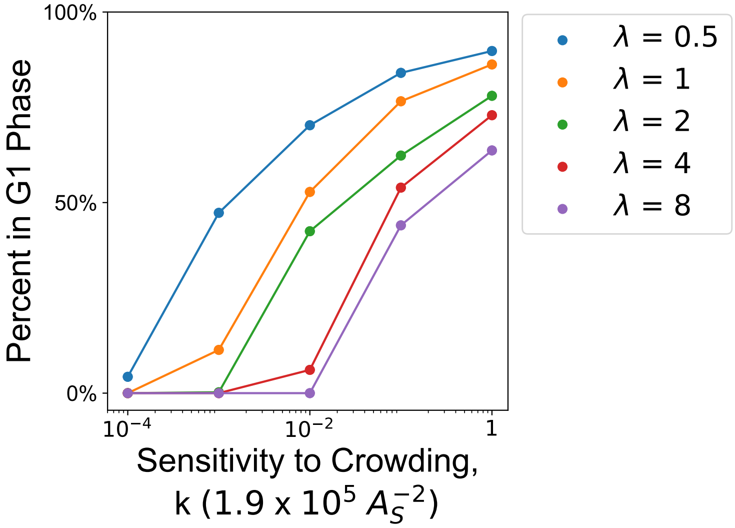


**Figure S** 3. Percentage of cells in the tissue in G1 phase, as a function of cellular sensitivity to crowding, $k$, for various values of cellular elastic modulus $\lambda$.
